# Socially regulated estrogen in an eavesdropping brood parasite

**DOI:** 10.1101/366799

**Authors:** Kathleen S. Lynch, Gulnoza Azieva, Anthony Pellicano

**Affiliations:** Hofstra University Hempstead NY 11549

**Keywords:** Brown-headed cowbird (Molothrus ater), social regulation, estrogen, hypothalamic-pituitary-gonadal axis (HPG)

## Abstract

Social regulation of reproductive hormones is a means by which conspecific males and females orchestrate successful reproductive efforts. There is variation, however, in the range of social cues that will initiate a hormone response in the receiver of social signals. We investigate whether social cues modify activity within the hypothalamic-pituitary-gonadal (HPG) axis and the specificity of this response in a social parasite that is known to eavesdrop on the communication signals of other species: the brown-headed cowbird (*Molothrus ater*). Brown-headed cowbirds are obligate brood parasites that do not build nests or care for their own young. Instead, obligate brood parasites always leave their eggs in the nest of a host species thereby receiving the benefits of parental care toward offspring without paying any of the costs. Thus, social parasites must coordinate their breeding attempts with conspecifics as well as potential heterospecific hosts and therefore, social parasites such as cowbirds rely on the communication signals of host species to help locate nests to parasitize during the breeding season. Here, we explore whether the vocal signals of potential host species can also be used as a social cue that modifies the HPG axis of female brown-headed cowbirds. Results reveal that both conspecific and heterospecific song-exposed females exhibit significantly greater circulating estradiol concentrations as compared to silence-exposed females. While conspecific song induces the greatest elevation in circulating estradiol, there is no significant difference in circulating estradiol levels in females exposed to either conspecific or heterospecific songs. This pattern suggests both song types are effective at evoking a reproductive physiological response. On the other hand, circulating progesterone concentrations did not differ among the song- and silence-exposed groups nor did the size of the female’s ovarian follicles. These results indicate that heterospecific vocal communication signals can effectively be used as a social cue that simultaneously provides necessary information regarding breeding status of hosts and modifies breeding condition of the eavesdropper.

Highlights

- Brood parasites may coordinate reproductive attempts with potential hosts.
- Therefore, parasites eavesdrop on other species, which helps them find nests.
- Vocal signals from conspecific and heterospecifics elevate estrogen in parasites.
- Hearing potential host songs may enhance reproductive hormones in brood parasites.

## 1.0 Introduction

Reproductive physiology and behavior can be regulated, in part, by social environment (Lehrman, 1965). Social regulation of reproductive physiology helps coordinate reproductive attempts between males and females that are attending to the social cues of their partners. There is variation, however, in the specificity of social signals that modify the receiver’s physiology. Reception of one’s own signals and/or reception of conspecific signals modify physiological states of the receiver, help coordinate complex social behaviors, and guide context-appropriate responses (Cheng, 1992; Cheng et al, 1998; Lynch and Wilczynski, 2006; Watts et al, 2016; Roleira et al, 2017). Here, we pair the phenomenon of social-dependent regulation of reproductive physiology with animal behavior studies demonstrating that some species intentionally and routinely eavesdrop on communication signals not intended for them. These species actively seek out signals for which they were not the intended target (i.e. eavesdropping) to obtain vital information that may enhance not just reproduction but also predation and parasitism opportunities, thereby increasing the eavesdropper’s fitness (Zuk and Kolluru, 1998; Bernal et al, 2006; 2007; Page and Ryan, 2008).

Social parasites that must locate hosts for reproductive purposes eavesdrop on the communication of heterospecifics. One example of a social parasite that eavesdrops is the brown-headed cowbird (*Molothrus ater*), an obligate brood parasite. Avian obligate brood parasites do not build nests, incubate eggs or provision their own young. Instead, the female parasite locates nests of other species that can serve as a host parent to her offspring. One of the ways in which breeding female parasites locate nests is through cryptic observation of hosts and their activities as well as attending to their vocalizations (Hann 1941; Gochfeld 1979; Clotfelter, 1998; Wiley, 1988; Alvarez 1993; Monk and Brush 2007; Janecka and Brush, 2014). This is termed the host-activity hypothesis, which predicts that brood parasites will be attracted to the vocalizations of potential hosts as well as their nest building behavior (Hann 1941; Gochfeld 1979; Wiley and Wiley 1980). Thus, conspicuous host vocalizations and other behaviors may be an important determinant of whether a host nest will be parasitized by eavesdropping brood parasites. Consequently, brood parasites should attend relevant social signals of conspecific and heterospecific species because the brood parasite must coordinate its reproductive behavior with both conspecific mates as well as possible heterospecific hosts.

The brown-headed cowbird (hereafter, cowbird) are oscine Passeriformes within the Icteridae family (i.e. blackbirds). Male cowbirds housed with females displayed peak circulating testosterone levels and mature gonads for longer as compared to cowbirds house without females (Dufty and Wingfield, 1990), indicating social regulation of reproductive physiology does occur in this species. Also, brown-headed cowbirds are known to eavesdrop on the activity of heterospecific hosts during the breeding season to aid in host nest location (Clotfelter, 1998). Here, we test the hypothesis that female cowbirds may coordinate their reproductive physiology with that of their hosts using social cues. We predict that heterospecific and conspecific vocalizations are both effective at modifying the reproductive physiology of female cowbirds. If this is the case, reproductive hormones and/or follicles will be significantly greater in conspecific and heterospecific song-exposed females as compared to silence-exposed females. These results will provide insight into whether brood parasites use heterospecific vocal signals as a means of coordinating timing of reproductive events and the specificity in the types of signals that will evoke a physiological response in these birds.

## 2.0 Methods

### 2.1 Housing and stimulus exposure

Female cowbirds (N = 18) were collected using bait traps in Travis and Kerr counties in Texas in May, 2016. Birds were transported to outdoor aviaries in Hempstead, NY and housed in semi-natural conditions. Females were fed a modified Bronx zoo diet with mealworm supplements and were isolated from male brown-head cowbirds to ensure no social interaction with males or male song during this time. In August, females showed no continued signs of breeding behavior. They were then housed in an indoor aviary to habituate for one week on a 16L:8D light schedule. Females were visually isolated from each other while individually housed in 610 mm × 610 mm cages randomly placed in indoor aviaries with varied social conditions: conspecific song exposure (N = 7), heterospecific song exposure (N = 7) or silence (N = 4). This design, including the sample sizes in each group, followed the design described in Bentley et al, (2000). All procedures presented here were permitted by Hofstra IACUC.

Red-winged blackbird (*Agelaius phoeniceus)* songs were chosen as the heterospecific stimulus in this study because they are a common host and geographically sympatric species for the brown-headed cowbird (Lowther, 1993). Red-winged blackbirds are frequently targeted as a host species for the brown-head cowbird and may even be the preferred species in some regions of the cowbirds range. Red-winged blackbirds have a similar breeding initiation timeframe as the cowbird, a lower nest abandonment rate and mite parasitism compared to other blackbirds (Freeman et al, 1990; Ortega and Cruz, 1988; 1990), all of which may result in the red-winged blackbird being a preferred host in some parts of North America. Furthermore, cowbirds are frequently found in mixed flocks that contain red-winged blackbirds. These observations indicate that red-winged blackbirds are likely to be encountered at some point in the cowbirds lifetime; either during developmental or adult stages. Stimuli were designed as described in Lynch et al., 2017. Briefly, five independent examples of song were recorded from different male brown-headed cowbirds (fig. 1a for representative sonogram) or red-winged blackbirds. Each sound was filtered above 2000 HZ and below 500 HZ, and all sounds were normalized to the mean amplitude. To equilibrate the amount of stimulation between experimental and control stimuli, we matched peak amplitude and duration of signals as described in Lynch et al, 2017. Vocalizations were synthesized with 20 s of vocal stimulus / min and arranged so that 1–2 songs from each male recorded was presented in each minute of presentation. The amplitude of song at each cage ranged between 65 and 70 dB as measured by SPL meter 0.5 m from the speaker. Songs were broadcasted for 8 hours / day starting at 6:00 a.m. followed by 16 hours of silence. Stimulus exposure in each condition continued for fourteen days during which time females continued to be fed the modified Bronx zoo diet with constant water supply. After fourteen days of song or silence exposure, females were sacrificed via rapid decapitation to collect blood tissue. As in Lynch et al (2017) and Bentley et al (2000), gonads were measured using calipers. The largest follicle was recorded in females across the three social exposure treatment groups.

**Figure 1.**
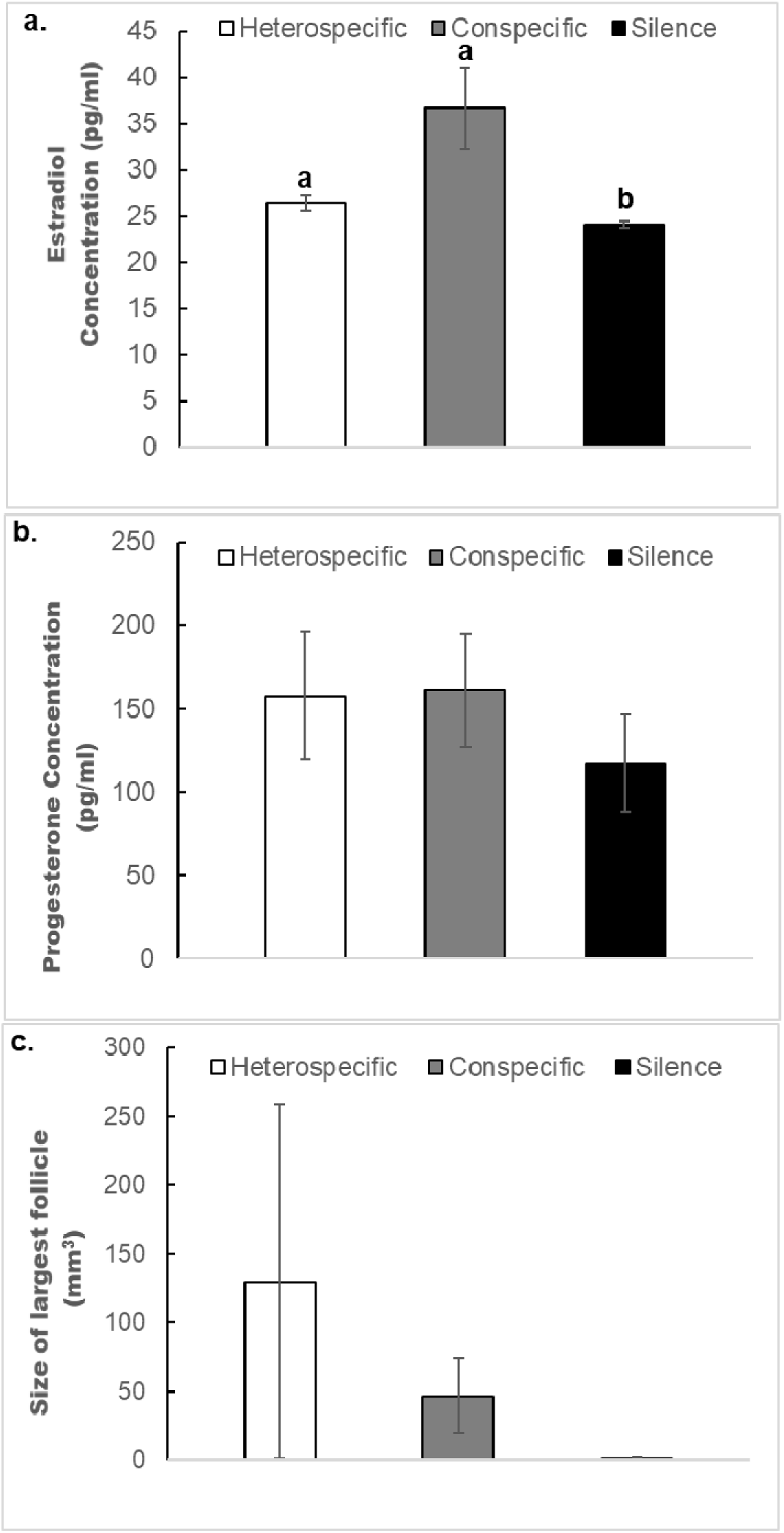
Results of hormone assays after female brown-headed cowbirds (*Molothrus ater*) were exposed to fourteen consecutive days of silence, conspecific or heterospecific songs. (a) Results comparing circulating estradiol assays, (b) Results comparing circulating progesterone assays and, (c) Results comparing female’s largest follicle. All error bars represent S.E.M.

### 2.2 Hormone assays

Immediately after collection, blood was centrifuged for 10min at 10,000rpm. Plasma was stored at −80°C until assayed. Circulating estradiol and progesterone concentrations were measured using modified extraction and assay procedures described in Lynch and Wilczynski, 2005; 2006; 2008. Briefly, steroids were extracted from the plasma using 3 ml of diethyl ether. Samples in all treatment groups were simultaneously extracted. In addition, a pooled sample of plasma was stripped of steroid using charcoal-dextran and spiked with a known concentration of estradiol (250 pg/ml) and measured in the EIA assay to estimate extraction efficiencies. The averaged extraction efficiency was calculated at 72%. Extracted steroids resuspended in assay buffer were used to measure estradiol and progesterone concentrations using ELISA assay kits from Cayman chemical (Ann Arbor, Michigan). To validate these kits for use with cowbird plasma, we extracted estradiol and progesterone from a plasma sample and serially diluted it at three concentrations. We compared the slope of the line for the serially diluted samples and the slope of the line for the area of the curve necessary to estimate the concentration of the diluted samples as described in Lynch and Wilczynski, 2006. The slope of the line for the serially diluted estradiol samples was −25.1, and the slope of the line for the area of the estradiol standard curve in which the samples are estimated was −27.9. The slope of the line for serially diluted progesterone samples was −811 and the slope of the line for the area of the progesterone standard curve in which samples are estimated was −792. Both steroids were measured on a single plate, precluding an inter-assay variation measurement. One subject was removed in the heterospecific group and two were removed from the conspecific group due to high CV estimates for the triplicates (final heterospecific N = 6; final conspecific N = 5). After removal of these two elevated CV samples, the intra-assay variation was 9.08% and 9.87% for estradiol and progesterone respectively. According to the manufacturer, estradiol EIA kits have a 0.1% cross reactivity with testosterone and 5 α-DHT, 0.07% for 17 α-estradiol, and 0.03% for progesterone and the detection limit is 6.6 pg/ml. The progesterone assay has a detection limit of 7.8 pg/ml and the following cross reactivities: pregnenolone 14%, 17β-estradiol 7.2%, 5β-pregnan-3α-ol-20-one 6.7%, and 17α-hydroxyprogesterone 3.6%. All other reported cross reactivities were less than 0.5%.

### 2.3 Statistics

Circulating estradiol and progesterone concentrations as well as follicle sizes were compared separately across three groups of females exposed to different social cues using non-parametric tests due to unequal variances. Separate Kruskall-Wallis tests determined whether there were any significant differences in circulating estradiol or progesterone concentrations across the three social exposure groups: conspecific and heterospecific song exposure as well as silence exposure. Post hoc pairwise comparisons were conducted using independent t-tests. Due to heteroscedasticity, pairwise tests were corrected for unequal variances. Benjamini-Hochberg corrections were used to adjust the false discovery rate for multiple tests as both conspecific and heterospecifc groups were compared to silence. This adjusted the heterospecific and conspecific comparison to silence to an alpha value of 0.025 and 0.05 respectively.

## 3.0 Results

A significant difference in circulating estradiol concentration across the three treatment groups was detected (H = 6.38; df = 2; P = 0.041). Post-hoc pairwise comparisons reveal that conspecific and heterospecific song exposed females had significantly higher circulating estradiol concentrations as compared to silence (t_4_= 2.83, P = 0.047; t_7_= 2.37, P = 0.022 respectively; fig. 1a). There was no difference in circulating estradiol concentrations between heterospecific and conspecific song exposed females (t_4_= −2.27, P = 0.08). On the other hand, no significant difference in circulating progesterone concentration across the three treatment groups was detected (H = 1.1; df = 2; P = 0.57; fig. 1b). Likewise, no significant difference was detected in largest follicle size (H = 2.23; df = 2; P = 0.32; fig. 1c).

## 4.0 Discussion

Social signals, including vocalizations, are sufficient to initiate activity within the hypothalamic-pituitary-gonadal (HPG) axis of birds (Bentley et al., 2000; Friedman, 1977; Kroodsma, 1976; Waas et al., 2005, Brockway, 1964; Haase et al., 1976). There is variation across species, however, in the specificity of social signals that will initiate this response (Bentley et al., 2000). In the present study, we apply this well-established phenomenon to the brown-headed cowbird, a species that eavesdrops on a wide array of heterospecific social signals to enhance its own fitness.

The results demonstrate that females exposed to either heterospecific or conspecific vocalizations displayed elevated circulating estradiol but not progesterone concentrations in comparison to females exposed to silence. These results indicate that both heterospecific and conspecific vocalizations will initiate a response in estradiol, but not progesterone, production in female cowbirds. Estradiol regulates a range of necessary processes in female songbirds during reproduction including modifying sensory processing that may enhance detection of male signals (Maney and Pinaud, 2011; Caras et al, 2012), inducing receptive responses to male courtship signals (Searcy and Capp, 1997; Leboucher et al, 1998) and, altering activity of catecholamines that also play a role in reproduction and mating (Matragrano et al, 2001; Rodríguez-Saltos et al, 2018; see Lynch, 2017 for review). Progesterone, on the other hand, has been demonstrated to inhibit reproductive behavior in some female songbirds (Leboucher et al, 2000). Thus, social regulation of circulating estradiol concentration by heterospecific and conspecific social signals may occur because both signals provide meaningful information regarding the commencement or conclusion of breeding season and allows brood parasitic birds to coordinate their reproductive efforts with each other as well as with their heterospecific hosts.

While circulating estradiol concentrations were significantly elevated by social exposure in female cowbirds, results indicate that development of ovarian follicles was unaffected by conspecific and heterospecific song exposure. There was no difference in measurements of either the largest follicle or the total size of the follicular cluster across any of the vocal exposure groups and silence. This result is consistent with studies from other songbirds, particularly pine siskins (*Spinus pinus*), European starlings (*Sternus vulgaris*), willow tits (*Parus montanus*) and white crowned sparrows (*Zonotrichia leucophrys pugetensis*). In these studies, females exhibited advanced gonadal development in the presence of a male, which appeared to be a vital cue instigating the progression to yolk deposition and follicular maturation (Perfito et al, 2014, Watts et al 2015; Wingfield et al., 1997). Therefore, it is possible that female cowbirds exposed to conspecific and heterospecific songs do not progress to advanced stages of follicular maturation because neither vocal signal is sufficient to initiate this process. As it is also the case that follicular development in song-exposed females is not different from females exposed to 14d of silence, it is possible that physical or visual interaction with males is necessary for advancing to mature follicular stages. This supports the hypothesis that vocal cues may be sufficient to instigate reproductive steroid production but final stages of follicle development require actual interaction with males (Watts et al 2015), suggesting that at least a visual cue from males are needed for the final stages of reproductive readiness to be achieved.

The results of this study reflect behavioral studies demonstrating that heterospecific songs provide meaningful information to brood parasites. For instance, in bronzed cowbirds (*Molothrus aeneus*), broadcasted songs of Audubon’s orioles are visited at nearly equal rates by orioles and bronzed cowbirds. Because orioles are a preferred host species for bronzed cowbirds this suggests that social cues used by orioles are a potent signal that attracts the attention of the cowbird (Monk and Brush, 2007). In a related study, bronzed cowbirds responded in greater numbers to the songs of orioles species (their preferred hosts) than to the songs of olive sparrows (*Arremonops rufivirgatus*), a lower-quality host species (Janecka and Brush, 2014). Further evidence indicates that female cowbirds use heterospecific songs as a cue during nest searching (Clotfelter, 1998; Janecka and Brush, 2014) and individual female cowbirds exhibit flexible preferences for a few specific host species (Strausberger and Ashley, 2005), which suggests memory for heterospecific signals. Thus, songs may help the brood parasite locate host nests and serve as a cue to help them find breeding heterospecifics that may soon build a nest. The results presented in our study suggest that signals from other species not only aid the brood parasite in locating host nests in which to lay eggs but also modulate the cowbirds reproductive hormones. Therefore, by attending to heterospecific signals the cowbird accomplishes two important tasks: it gains information regarding the possible location of nests while simultaneously coordinating its reproductive timing with that of its host species.

Social regulation of reproductive physiology via heterospecific signals has occurred in other species albeit not to the extent that conspecific signals evoke the response. For instance, in captive canaries, conspecific song exposure induced marginally significantly greater follicle sizes as compared to heterospecific song exposure but both social exposure groups exhibited significantly greater follicular sizes in relation to silence-exposed females (Bentley et al., 2000). However, when time to lay eggs was measured, conspecific song-exposed females displayed significantly shorter time to lay eggs and laid more of them in relation to heterospecific song-exposed females. Taken together, these experiments reveal conspecific song-exposed female exhibited the greatest responses in ovarian development however, heterospecific song exposure induced a greater response in comparison silence exposure in these measures. This suggests that even in a captive population of canaries, heterospecific songs possess some ecological relevance, albeit less than conspecific songs. This result implies that, to some degree, all birds attend to communication signals not intended for them and that these signals possess ecological relevance as they may inform the receiver of major transitions within the avian community regarding the breeding or non-breeding status of the members of the community.

From an evolutionary perspective, physiological responses to song are likely subjected to less selection pressure for discrimination than the female’s behavioral response. Thus, a broader range of signals should evoke activity within the HPG axis while only a subset of the most relevant signals should evoke a behavioral response from a female. By broadening the definition of signals that can evoke physiological responses, animals can orchestrate reproductive efforts across multiple biological levels including between individuals, populations and communities. Consequently, less attractive conspecific signals and even heterospecific signals may evoke physiological responses in reproductive systems in eavesdropping animals and possibly in wide array of free-living animals.

## Acknowledgments

We thank Texas EcoLabs and Balcones Wildlife Refuge for support with field collections of cowbirds and financial support of this study. We also thank Mary Ramsey for comments on earlier versions of this manuscript.

